# Negative relationship between inter-regional interaction and regional power: a resting fMRI study

**DOI:** 10.1101/2024.01.31.578128

**Authors:** Tien-Wen Lee, Gerald Tramontano

## Abstract

**Background:** Regional neural response and network property used to be treated separately. However, evidence has suggested an intimate relationship between the regional and inter-regional profiles. This research aimed to investigate the influence of functional connectivity on regional spontaneous activity.

**Methods:** Thirty-six and sixty datasets of structural magnetic resonance imaging (sMRI) and resting state functional MRI (rsfMRI) were selected from the NKI and CAN-BIND database, respectively. The cerebral cortex in rsfMRI was parcellated by MOSI (modular analysis and similarity measurements), which enables multi-resolution exploration. For each parcellated cluster, the mean amplitude of low-frequency fluctuation (ALFF) and its average functional connectivity strength with the remaining cortical analogs were computed. Correlation analyses were exploited to examine their relationship. Supplementary analysis was applied to CAN-BIND EEG data (1 to 30 Hz).

**Results:** Negative correlation coefficients between inter-regional interaction and regional power were noticed in both MRI datasets. One-sample t-tests revealed robust statistics across different analytic resolutions yielded by MOSI, with individual *P* values at the level 10^-4 to 10^-5. The results suggested that the more intense crosstalk a neural node is embedded in, the less regional power it manifests, and vice versa. The negative relationship was replicated in EEG analysis but limited to delta (1 to 4 Hz) and theta (4 to 8 Hz) frequencies.

**Conclusions:** We postulate that inhibitory coupling is the mechanism that bridges the local and inter-regional properties, which is more prominent in the lower spectra. The interpretation warrants particular caution since noise may also contribute to the observation.

## Introduction

Regional brain activity is embedded within a context of intense crosstalk mediated by vast neural interactions. Previous neuroimaging research has typically investigated regional neural activity or inter-regional connectivity separately. An earlier electroencephalography (EEG) study reported that regional entropy was predicted by the centrality index, the measure of a node’s relative importance within a graph (Misic et al., 2011). Our previous work showed that the degree of global network synchronization in EEG, especially at the alpha spectrum, was associated with higher neural responsiveness and faster neural propagation in early evoked components (Lee et al., 2011). In another study with magnetic resonance imaging (MRI) we highlighted that regional amplitude of low-frequency fluctuation (ALFF) of functional MRI (fMRI) signals was negatively correlated by the sum of its structural connectivity, as indexed by diffusion tensor imaging (DTI) (Lee and Xue, 2017). Together, these findings suggest a relationship between regional and inter-regional neural metrics.

Previous rsfMRI studies demonstrated a complicated and less than inconsistent relationship between ALFF and functional connectivity (FC) within selected networks (Di et al., 2013; Gan et al., 2022). For example, the relationships between ALFF and FC within a functional network characterized by independent component analysis are generally positive (see Figure 4 in (Di et al., 2013)). The default-mode network component was positively and negatively correlated with the ALFF in the posterior cingulate cortex and inferior parietal region, respectively (see Figure 1 in (Di et al., 2013)). On the other hand, mindfulness training was reported to drive the ALFF and FC within the saliency network in opposite directions (Gan et al., 2022). It is justifiable to examine these neural metrics within a network; however, given the broad and distributional interconnections of the brain, it is equally important to examine the issue at a global scale (Deco et al., 2011; Lee, 2016; Lee et al., 2014; van den Heuvel and Hulshoff Pol, 2010; Xue et al., 2014), as illustrated above in (Lee and Xue, 2017). To the best of our knowledge, whether or how the overall FC profile of a neural node may be inter-related with its regional power, as indicated by ALFF, has never been explored.

**Figure 1.**
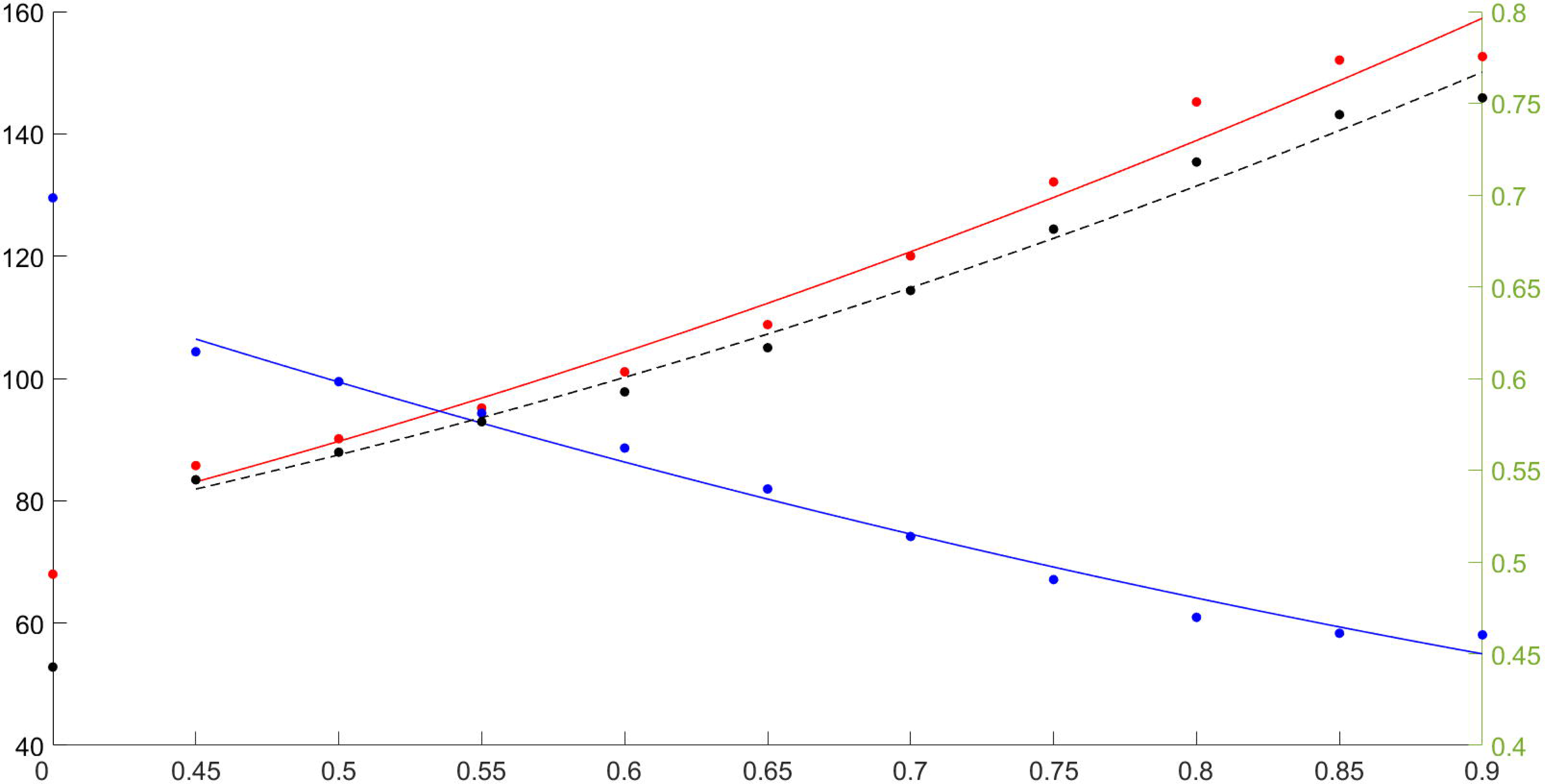
Scatter plot of three indices derived from NKI dataset, with X-axis gamma values and second-order polynomial regression fits. The origin (x = 0) represents the initial partition based on the Desikan-Killiany atlas. Red: mean module number of the partitions derived from different gamma values, referring to the left Y-axis. Blue: mean voxel number per module, referring to the left Y-axis. Black: mean module z-score (homogeneity within a module), referring to the right Y-axis.

Although the issue looks straightforward, an innate difficulty must be overcome to enable the assessment. Current fMRI may record neural information at a spatial resolution of several millimeters, allowing the brain to be divided into tens to hundreds of thousands of voxels. While enhanced resolution is a favorable development, the abundance of voxels presents an obstacle in analyzing brain networks. As the temporal dynamics between neighboring fMRI voxels are not independent, investigators have applied dimension reduction strategies to simplify the data structure. One common approach for cortical partition is to divide the brain into distinct regions based on anatomy, although its validity has frequently been questioned (Amunts et al., 1999; Lee et al., 2014; Smith et al., 2011; Stanley et al., 2013; Wig et al., 2011; Xue et al., 2014). It would be ideal to partition the brain dynamics based on fMRI signal itself, i.e., functional parcellation (FP), in contrast to relying on the atlas-informed structural parcellation (Damoiseaux et al., 2006; Shen et al., 2013; Shi and Malik, 2000; Smith et al., 2009). The authors developed an innovative analytic scheme – modular analysis and similarity measurements (MOSI) – to address this issue (detailed in **Methods**) (Lee and Tramontano, 2021b). A data-driven algorithm, MOSI requires no extra premises and can substantiate individualized and multi-resolution parcellations.

In this study, with resting-state fMRI (rsfMRI) data of two publicly released datasets, we aimed to adopt MOSI to segment the cortex into functionally homogeneous clusters. Each cluster would be composed of voxels with similar traces of BOLD signals and hence designated as a neural node. The constituent nodes “in communication” defines a network. The ALFF was computed for each node, while the inter-nodal interaction was represented by the degree of correlation between the paired mean time series, i.e., FC. The relationship between the regional and inter-regional profiles can thus be examined systematically across different spatial resolutions. Based on our previous cross-modal study that the ALFF (rsfMRI) decreases with nodal degree (DTI) (Lee and Xue, 2017), we hypothesize that the network substantiated via inter-nodal interaction will shape regional activity in rsfMRI; specifically, the overall FC of a node and its regional power would show a negative correlation.

## Material and methods

### Subjects, MRI data, and preprocessing

We employed the data of a total of 96 healthy subjects, including 36 (average age 27.4 years) from the Rockland sample dataset and 60 (average age 33.2) from the Canadian Biomarker Integration Network in Depression (CAN-BIND) (Lam et al., 2016; Tobe et al., 2022), respectively. The functional and structural MRI (sMRI) images of the entire brain were recorded using 3.0 Tesla MRI machines (NKI: Siemens, TR=2.5s, voxel size=3×;3×3mm^3, 255 volumes; CAN-BIND: GE, TR=2s, voxel size=4×4×4mm^3, 300 volumes). For more information about the scanning protocol, please visit the webpages http://fcon_1000.projects.nitrc.org/indi/pro/nki.html and the study protocol in (Lam et al., 2016). The two datasets were analyzed separately.

We processed the echo planar imaging (EPI) data using the Analysis of Functional NeuroImages software package (AFNI) (Cox, 1996). The preprocessing steps of rsfMRI comprised despiking, slice-time correction, realignment (motion correction), registration of T1 anatomy, spatial smoothing, and bandpass filtering at 0.01–0.1 Hz. The first 4 scans were disregarded. Smoothness kernels 6 mm and 8 mm were respectively applied to NKI and CAN-BIND EPI data. The software FreeSurfer was adopted to segment gray and white matters from T1-weighted sMRI data and to divide the cortical mantle into 68 regions of interest (ROIs; Desikan-Killiany Atlas) (Dale et al., 1999; Desikan et al., 2006; Fischl et al., 1999), which provided the initial partition for subsequent MOSI analysis. Following the analytic pipeline developed by Jo et al. (Jo et al., 2010; Lee et al., 2014; Xue et al., 2014), 12 movement parameters, and white matter (both global and local) and ventricular signals were modeled in the regression (Jo et al., 2010). Further, third-order polynomials were applied to fit baseline drift. ALFF was computed as the square root of the band-passed power of BOLD dynamics.

### Functional connectivity and MOSI analysis

Pearson correlation coefficient (CC) was computed pairwise between the mean temporal traces of any paired voxels or clusters to indicate FC. The connectivity map based on CCs is the foundation for modular analysis. Modules or communities are groups of nodes within a network that are more densely connected to one another than to other nodes, as defined by the CCs. MOSI incorporates modular analysis and similarity measurements, splitting a module into sub-modules and uniting similar sub-modules into a bigger module. MOSI fulfills the parcellation of the cortex abiding by only two criteria: neighborhood and similarity, with adjacent voxels sharing similar neural activities grouped together, precisely the rationale behind FP of cortical signals. The splitting-unifying processes were repeated until convergence. The current version of MOSI uses the Louvain community detection algorithm. The number of modules and, hence, the partition resolution can be titrated by gamma values, with a higher value yielding more modules (Lee and Tramontano, 2021b). We explored gamma values ranging from 0.45 to 0.90 (10 different resolutions). The unique features of MOSI include: (1) it is an individualized analysis; (2) it lays out a plausible foundation for subsequent network analysis; (3) it enables a multi-resolution approach to investigate brain informatics at different scales (Lee and Tramontano, 2021b).

At the single subject level, each module’s mean ALFF and z-score (the CCs were converted to z-scores by Fishers’ transformation) were calculated for each partition, for each gamma value. Assuming a module has N voxels, the average “module z-score” was derived by taking an average of N*(N-1)/2 of all possible directionless connections’ z-scores (of CCs) within that module, and the mean ALFF of the N voxels was computed for that module. Assuming that MOSI output M modules at a particular gamma value (partition), a correlation between the mean ALFFs and module z-scores was computed across all derived M modules, resulting in 10×36 (NKI) plus 10×60 (CAN-BIND) correlation analyses in total. Each correlation was associated with a *P* value that will help yield an average individual-level *P* value by taking the geometrical mean (see below), which were imported to group-level analyses after another Fisher’s transformation (partition z-score). Note that the “module z-score” was positive given that the BOLD signals were similar within a module and thus positively correlated, whereas the “partition z-score” was expected to be negative, based on our prediction that the ALFFs and module z-scores (CCs) were negatively correlated.

### Statistical analyses

For each gamma value and the associated partition, a one-sample t-test was performed with a null hypothesis that the partition z-score equaled zero – no correlation between ALFF and connectivity strength. The statistical threshold was set at *P* < 0.0025 to correct for multiple comparisons (0.05/20, with ten gamma values for two datasets). Since at the individual level, each partition z-score was accompanied by a *P* value, we used geometric mean to derive a representative individual-level *P* value: There were 36 subjects from NKI sample, with the *P* values of correlation p1, p2, …, and p36 for each partition. The representative *P* was calculated as (p1*p2*…pk)^(1/36). A similar computation was applied to the 60 CAN-BIND subjects.

### Supplementary analyses

In our previous study that explored the relationship between ALFF and structural connectivity, the authors premised that the negative relationship could be mediated by the dissimilarity of the inputs to a neural node (receiver). Since the neural dynamics are complicated if the interactions between a neural node and the remaining ones were effective enough, there is a chance that the peaks and troughs of the incoming signals may cancel each other and their influence on the receiver node would diminish—higher connectivity causing lower input and hence lower regional power reflected in the receiver node (Lee and Xue, 2017). To examine the conjecture, we introduced two indices as regressors. We computed first, the average correlation strengths among the inputs (after Fisher’s transformation) and second, the average mutual information between the inputs (the bin numbers were determined by Sturge’s Rule, ten for NKI and eleven for CAN-BIND (Sturges, 1926)). The former and latter respectively addressed the temporal and distributional inter-dependence.

Since BOLD fluctuation in rsfMRI approximates low-pass filtered neural activity (< 0.1 Hz), we resorted to an EEG dataset from CAN-BIND to explore the power–connectivity issue at higher spectra, where we focused on delta to beta range (1–30 Hz). CAN-BIND provides 64-channel EEG data (58 channels after harmonization) from healthy controls. We employed the software EEGLAB to edit the EEG traces (Delorme and Makeig, 2004). Data preprocessing included band-pass filtering (1–50 Hz), automatic artifact removal (Artifact Subspace Reconstruction), and manual elimination of the remaining noisy portions. A single rater (TW Lee) carried out the above processing. Quality screening left 47 out of the 54 controls in the analyses. With the artifacts, including those associated with blinks and eye movements, removed, the cleaned EEG data were segmented into 2-sec epochs and imported to eLORETA for subsequent analyses (Pascual-Marqui, 2007a). With eLORETA, the neural informatics from the electrodes can be projected to a Talairach brain template with 6,239 gray matter voxels. We then retrieve the mean power of and the connectivity strength between the 68 cortical regions defined by, again, Desikan-Killiany atlas (Desikan et al., 2006).

## Results

Concordant with the nature of MOSI, the cluster number increased, and the number of voxels in each cluster decreased with the gamma values. For NKI sample, the segmented module numbers of the cortex were 97.3 and 183.9 at gamma 0.45 and 0.90, respectively, with the module z-scores increasing from 0.55 (CC 0.50) to 0.76 (CC 0.64), see Figure 1. The module number of the initial partition based on the Desikan-Killiany Atlas was 68, with an average voxel number of 253.2 (27mm^3/voxel) and a module z-score of 0.40 (CC 0.38). For the CAN-BIND sample, the segmented module numbers of the cortex were 85.8 and 152.6 at gamma 0.45 and 0.90, respectively, with the module z-scores increasing from 0.54 (CC 0.49) to 0.75 (CC 0.64), see Figure 2. The module number of the initial partition based on the Desikan-Killiany Atlas was 68, with an average voxel number of 129.5 (64mm^3/voxel) and a module z-score of 0.44 (CC 0.41).

**Figure 2.**
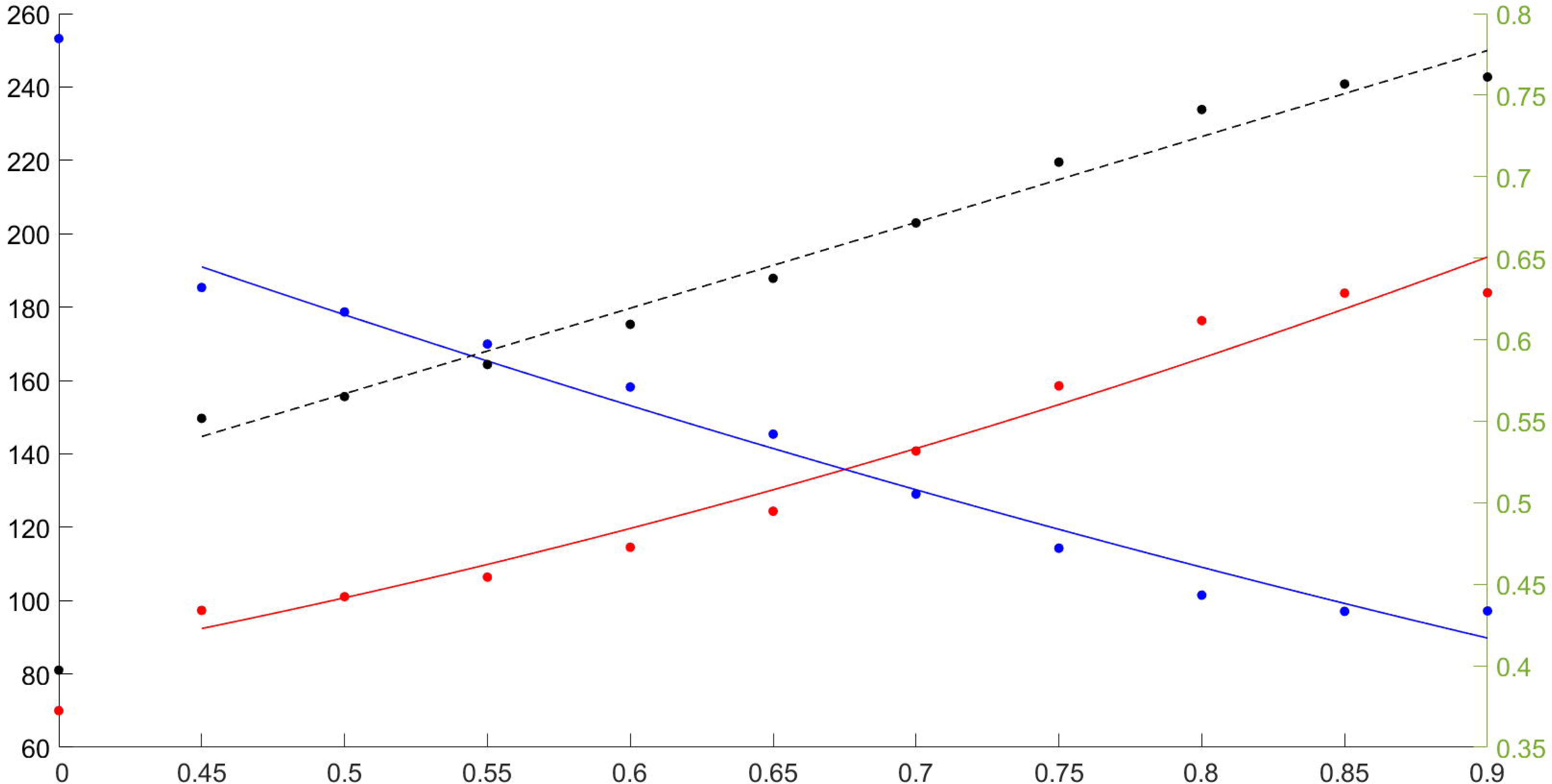
Scatter plot of three indices derived from CAN-BIND dataset, with X-axis gamma values and second-order polynomial regression fits. The origin (x = 0) represents the initial partition based on the Desikan-Killiany atlas. Red: mean module number of the partitions derived from different gamma values, referring to the left Y-axis. Blue: mean voxel number per module, referring to the left Y-axis. Black: mean module z-score (homogeneity within a module), referring to the right Y-axis.

As predicted, the module z-scores and ALFFs showed robust negative correlation across all the different resolutions for both datasets. The representative individual *P* value ranged from 7.19E-04 to 4.07E-05 for NKI and 1.22E-07 to 5.65E-08 for CAN-BIND samples. Regarding partition z-scores for NKI data, the group-level t-statistics and *P* values of the one-sample t-tests ranged from −8.77 to −11.80 and 2.32E-10 to 9.39E-14, respectively, see Table 1. As for partition z-scores for CAN-BIND data, the group-level t-statistics and *P* values of the one-sample t-tests ranged from −23.41 to −27.16 and 1.50E-31 to 4.81E-35, respectively, see Table 2.

**Table 1.** Statistics of the correlation between ALFF and functional connectivity strength across different gamma values (from NKI)

| Gamma | 0.45 | 0.50 | 0.55 | 0.60 | 0.65 | 0.70 | 0.75 | 0.80 | 0.85 | 0.90 |
| --- | --- | --- | --- | --- | --- | --- | --- | --- | --- | --- |
| CC | -0.37 | -0.36 | -0.34 | -0.34 | -0.32 | -0.30 | -0.28 | -0.25 | -0.21 | -0.21 |
| Mean_Z | -0.36 | -0.34 | -0.33 | -0.32 | -0.31 | -0.29 | -0.27 | -0.24 | -0.21 | -0.21 |
| Std_Z | 0.18 | 0.18 | 0.18 | 0.18 | 0.17 | 0.16 | 0.15 | 0.14 | 0.14 | 0.14 |
| t stat | -11.80 | -11.34 | -10.91 | -10.93 | -10.70 | -10.70 | -10.84 | -10.59 | -8.77 | -8.86 |
| P value | All *P* values are less than 0.005, ranging from 2.32E-10 to 9.39E-14. |  |  |  |  |  |  |  |  |  |
| Pg value | 4.07E-05 | 5.63E-05 | 7.49E-05 | 5.27E-05 | 6.60E-05 | 6.94E-05 | 1.09E-04 | 1.98E-04 | 6.37E-4 | 7.19E-4 |

Note: CC is the correlation coefficient converted from the mean z-score (Mean\_Z) by the function tanh. Pg is the geometrical mean of the *P* values derived from all subjects (N=36).

**Table 2.** Statistics of the correlation between ALFF and functional connectivity strength across different gamma values (from BrainCode)

| Gamma | 0.45 | 0.50 | 0.55 | 0.60 | 0.65 | 0.70 | 0.75 | 0.80 | 0.85 | 0.90 |
| --- | --- | --- | --- | --- | --- | --- | --- | --- | --- | --- |
| CC | -0.42 | -0.41 | -0.41 | -0.41 | -0.40 | -0.39 | -0.38 | -0.35 | -0.33 | -0.32 |
| Mean_Z | -0.45 | -0.44 | -0.43 | -0.44 | -0.42 | -0.41 | -0.40 | -0.37 | -0.34 | -0.33 |
| Std_Z | 0.19 | 0.19 | 0.18 | 0.18 | 0.18 | 0.16 | 0.15 | 0.14 | 0.14 | 0.15 |
| t stat | -27.16 | -26.63 | -25.70 | -26.01 | -23.87 | -25.79 | -23.85 | -24.30 | -23.41 | -24.46 |
| P value | All *P* values are less than 0.005, ranging from 1.11E-24 to 5.20E-30. |  |  |  |  |  |  |  |  |  |
| Pg value | 2.53E-05 | 2.52E-05 | 1.70E-05 | 8.76E-06 | 6.96E-06 | 4.18E-06 | 3.14E-06 | 4.39E-06 | 1.05E-5 | 1.57E-5 |

Note: CC is the correlation coefficient converted from the mean z-score (Mean\_Z) by the function tanh. Pg is the geometrical mean of the *P* values derived from all subjects (N=60).

### Results of supplementary analysis

As for fMRI data, after controlling the influence of the two regressors, the negative relationships between functional connectivity and regional power are still robust, as summarized in Supp Tables 1 and 2. As for EEG data, it is interesting to note that the inverse relationship between regional power and inter-regional FC holds for delta and theta spectra, but not alpha and theta counterparts, see Supp Table 3.

## Discussion

A neural network can be defined by two classes of descriptors, i.e., nodal properties and how the composite neural nodes interact. A large portion of empirical research explored regional neural activities and inter-regional structures separately. Nevertheless, evidence has suggested that the context in which a neural node resides may shape its activity (Lee and Xue, 2017; Lee et al., 2011; Misic et al., 2011). We extended previous research and examined the impact of mean FC on regional property ALFF. MOSI was adopted to achieve FP and a multi-resolution intervention. It was shown that nodal ALFF decreased with average inter-nodal interactions, and the statistics were robust. The group-level t-statistics indicated a consistent trend across subjects, with geometric P-values determining the significance of the trend for each individual. Our hypothesis was confirmed. In addition, the module z-scores were better than atlas-information counterparts and monotonously increased with gamma values, see Figures 1 and 2, indicating that the BOLD signals within a module were more homogeneous with finer resolution, which verified the validity of the MOSI scheme. The results from both NKI and CAN-BIND agree with the previous methodology report by the authors that MOSI may reliably achieve FP of cortical signals without extra constraints or assumptions. It is interesting to note that our supplementary analysis of the EEG dataset also replicated the findings at a lower frequency range (1–8 Hz; supplementary II). Considering the authors’ previous study that enhanced structural connectivity was accompanied by decreased regional ALFF, it is tenable to conclude that regional power and global inter-regional crosstalk, either functional or structural, may proceed in opposite directions. Moreover, it is more appropriate to place the power shifts in the context of discussions on changes in connectivity (assuming global structural connectivity strength is the determinant of local power change, not vice versa), with an awareness that correlation does not imply causation (Lee, 2017 #20948). Specifically, we will interpret the discovery as the overall network influence (average functional connectivity to a node) on regional power, rather than the other way around. Then, what are the mechanisms underlying this robust Inverse interaction?

The primary factor that should be considered is noise. Assume there are independent noises added to two neural nodes. The power of the two nodes will hence increase and given the independence of the introduced randomness, their connectivity strength will decrease. In fMRI, movement may introduce synchronized and systemic noise, which is expected to reflect in the enhanced connectivity, and because of the dephasing of the small proton magnets, the BOLD signal intensity is expected to attenuate (Power et al., 2012; Van Dijk et al., 2012; Yan et al., 2013). A full discussion about other physiological and non-physiological MRI artifacts will not be covered here (Birn, 2012; Power et al., 2014). Although routine measures have been adopted in the pre-processing of fMRI data to protect the BOLD signals from the contamination of noise, the robust statistics are inevitably, at least partly, contributed by noise. However, noise is unlikely to be the only cause of the findings. The previous report of structural connectivity and ALFF cannot be explained simply by noise (Lee and Xue, 2017). Moreover, a negative correlation implies a linear relationship, which is hard to reconcile by regional or systemic noise. For example, the head movement needs to “design” itself to simultaneously attenuate the signals in neural nodes N1, N2, N3 … to the degree that their FC with the remaining nodes shows opposite and proportional increments. Noise may accommodate the observation in a simple scenario such as a two-node situation as exemplified above, but is difficult to account for the observed group-to-one relationship (i.e., averaged FC as group-to-nodal property). If noise is the major determinant of the findings, then negative correlation implies that the region suffering strongest signal loss in the EPI image would have the highest correlation with the whole brain—a very unlikely possibility.

This study was also inspired by an anecdotal observation. We noticed that neuropsychiatric patients who demonstrated widespread decreased connectivity (using the in-house NeuroStation platform) were usually accompanied by abnormally increased regional ALFF. Our initial interpretation resorted to a compensatory mechanism that local neural tissue needs to work harder to compensate for the inadequacy of inputs from the surroundings. However, this research data was obtained from healthy volunteers, so the discovery demands alternative interpretations. Putting aside noise, we formulated two conjectures to account for this phenomenon.

Firstly, given the complexity of neural dynamics, a brain region with higher connectivity to other brain regions is expected to receive more diverse neural inputs, and hence the similarity between the BOLD dynamics of its composite voxels is expected to be lower. Based on the above inference, we surmised that diverse inputs might render the BOLD dynamics of the voxels in a brain region variable and out-of-phase, which is expected to cancel off each other after smoothing and hence reduce the mean ALFF (explicated in (Lee and Xue, 2017)). This assumption is reasonable but mundane and is not supported by our extra analysis (*supplementary material I*). It is important to consider that the dynamics of “regional” neural tissue itself are complex and have innate variability and chaotic features in which small input differences can lead to significant changes in the resulting manifestation (Chialvo, 2010; Freeman et al., 1997; Lee, 2016; Lee and Tramontano, 2021a). The above interpretation as an explanatory candidate is thus excluded.

Our second conjecture proposes that the lower connectivity in rsfMRI is associated with less neural inhibition instigated from the inputs of neighboring or distant neural nodes, thus enhancing regional power. Furthermore, lower functional connectivity indicates, and may result from, less organized inhibitory engagement and in turn renders the two regions’ dynamics less similar. This interpretation accords with the neuroscientific understanding that inhibitory coupling is critical in synchronizing the neural dynamics of networks (Andreev et al., 2021; Reimbayev et al., 2017). Similar analyses were applied to EEG data (*supplementary material II*). The results showed an interesting pattern that at lower frequencies (delta and theta), there remained a robust negative relationship. At a higher frequency range (alpha to beta), the negative relationship disappeared. In EEG, unlike rsfMRI, artifacts may increase regional power and inter-regional connectivity concurrently, not oppositely, through say, volume conduction or simultaneous involvement of several electrodes (Pascual-Marqui, 2007b). BOLD fluctuation in rsfMRI approximates low-pass filtered neural activity (< 0.1 Hz) (Lee and Xue, 2018; Xue et al., 2014). Together with the finding of spectrum-specificity in EEG, we argue that the information transmitted at a lower frequency range (fMRI and EEG; up to 8 Hz) may play various roles related to inhibitory interaction.

For the neuropsychiatric population, extra mechanisms may contribute to the observed negative relationship between inter-regional connectivity and regional power. It is known that around 75% of neocortical neurons are excitatory, while inhibitory interneurons comprise the rest one-fourth. To maintain network equilibrium, the firing rates and synaptic strength of inhibitory interneurons are higher than those in excitatory neurons, and the depression of inhibitory synapses due to sustained activation is less significant (see an excellent review by (Markram et al., 2004)). The inhibitory organization is more vulnerable to structural damage than the excitatory counterpart, with the latter equipped with a more abundant reserve. The proposition may also accommodate the previous report on structural connectivity and ALFF (Lee and Xue, 2017). In summary, the observed negative relationship between regional and inter-regional neural informatics is multi-factorially determined and may unveil coupled neural inhibitory control.

## Conclusion

Previous cross-modal MRI research and anecdotal clinical cases disclosed that regional power and its degree of interaction with other brain areas were negatively correlated. This study applied MOSI to rsfMRI data and replicated the discovery. The relationship was consistently observed across different resolutions of MOSI, statistically robust, and replicated in the delta and theta range of EEG. The findings altogether point to a fundamental brain principle that bridges the local and inter-regional properties and is most likely mediated by inhibitory coupling at a lower frequency range.

## Authors Contributions

Both authors contributed intellectually to this work. TW Lee carried out the analysis and wrote the first draft. Both authors revised and approved the final version of the manuscript.

## Acknowledgments

We want to thank Enhanced Nathan Kline Institute-Rockland Sample (NKI-RS) and Canadian Biomarker Integration Network in Depression (CAN-BIND) for their generosity in sharing MRI and EEG data. This work was supported by NeuroCognitive Institute (NCI) and NCI Clinical Research Foundation Inc.

## Statements and Declarations

Both authors declare no conflicts of interest.

## Compliance with ethical standards

This research analyzed the databank from a publicly released dataset. The authors carried out no animal or human studies for this article.

