## Supplementary Material for "Negative relationship between inter-regional interaction and regional power: a resting fMRI study"

**Supplementary Materials**

The **supplementary materials** comprise two parts. The first part is the re-analyses of the fMRI data with two extra regressors. The second part explores the same issue using a resting EEG dataset.

I. **Re-analyses of the fMRI data**

It was speculated that the negative relationship between connectivity and power could be mediated by the (dis)similarity between inputs to a neural node (receiver). If the inputs were dissimilar, e.g., because of the complicated neural dynamics and their interactions with that neural node were effective enough, there is a chance that the peaks and troughs of the incoming signals may cancel each other and their influence on the receiver node would diminish—higher connectivity causing lower input and hence lower regional power reflected in the receiver node (1). We introduced two indices as regressors to decipher this issue. We computed first, the average correlation strengths among the inputs (after Fisher’s transformation) and second, the average mutual information between the inputs (the bin numbers were determined by Sturge’s Rule, ten for NKI and eleven for CAN-BIND (2)). The former and latter respectively addressed the temporal and distributional inter-dependence. After partialling out the influence of the two regressors, the negative relationships between functional connectivity and regional power are still robust, summarized in **Supp Tables 1** and **2**.

**Supp Table 1**

Statistics of the correlation between ALFF and functional connectivity strength across different gamma values (from NKI)

Gamma	0.45	0.50	0.55	0.60	0.65	0.70	0.75	0.80	0.85	0.90

CC	-0.34	-0.32	-0.30	-0.31	-0.28	-0.27	-0.25	-0.22	-0.19	-0.18

Mean_Z	-0.35	-0.33	-0.31	-0.32	-0.28	-0.28	-0.25	-0.23	-0.19	-0.18

Std_Z	0.22	0.22	0.23	0.19	0.17	0.18	0.15	0.14	0.14	0.15

t stat	-9.47	-9.08	-8.12	-10.09	-9.72	-9.36	-9.73	-9.46	-8.38	-7.45

*P* value	All *P values are* less than 0.005, ranging from 1.01E-8 to 6.74E-12.

*Pg* value	2.03E-04	3.15E-04	3.55E-04	2.31E-04	5.97E-04	3.07E-04	6.73E-04	9.18E-04	1.86E-3	2.68E-3

Note: CC is the correlation coefficient converted from the mean z-score (Mean_Z) by the function tanh. Pg is the geometrical mean of the *P* values derived from all subjects (N=36).

**Supp Table 2**

Statistics of the correlation between ALFF and functional connectivity strength across different gamma values (from BrainCode)

Gamma	0.45	0.50	0.55	0.60	0.65	0.70	0.75	0.80	0.85	0.90

CC	-0.40	-0.38	-0.38	-0.39	-0.38	-0.36	-0.36	-0.35	-0.31	-0.29

Mean_Z	-0.43	-0.40	-0.40	-0.41	-0.41	-0.38	-0.38	-0.36	-0.32	-0.30

Std_Z	0.18	0.18	0.17	0.17	0.17	0.16	0.14	0.14	0.13	0.14

t stat	-18.67	-17.45	-18.71	-18.72	-18.37	-18.07	-20.96	-19.84	-18.62	-17.08

*P* value	All *P values are* less than 0.005, ranging from 1.61E-24 to 5.03E-29.

*Pg* value	6.13E-05	9.53E-05	6.21E-05	3.01E-05	1.47E-05	2.01E-05	7.38E-06	6.19E-06	3.50E-5	7.28E-5

Note: CC is the correlation coefficient converted from the mean z-score (Mean_Z) by the function tanh. Pg is the geometrical mean of the *P* values derived from all subjects (N=60).

II. **Exploration of power—connectivity relationship using EEG data**

Since BOLD fluctuation in rsfMRI approximates low-pass filtered neural activity (< 0.1 Hz), we resorted to an EEG dataset from CAN-BIND to explore the power–connectivity issue at higher spectra, where we focused on delta to beta range (1–30 Hz). CAN-BIND provides 64-channel EEG data (58 channels after harmonization) from healthy controls. We employed the software EEGLAB to edit the EEG traces (3). Data preprocessing included band-pass filtering (1–50 Hz), automatic artifact removal (Artifact Subspace Reconstruction), and manual elimination of the remaining noisy portions. A single rater (TW Lee) carried out the above processing. Quality screening left 47 out of the 54 controls in the analyses. With the artifacts, including those associated with blinks and eye movements, removed, the cleaned EEG data were segmented into 2-sec epochs and imported to eLORETA for subsequent analyses (4). Based on the principles of linearity and superposition, eLORETA is suitable for delineating distributed electric sources (current source density; CSD) in the brain cortex, albeit with low spatial resolution. With eLORETA, the neural informatics from the electrodes can be projected to a Talairach brain template with 6,239 gray matter voxels. We then projected the CSD time series to 70 cortical regions by Desikan-Killiany atlas (5). Average time courses derived for each region. The power spectrum of delta (1–4 Hz), theta (4–8 Hz), alpha1 (8–10 Hz), alpha2 (10–12 Hz), alpha (8–12 Hz), low beta (12–18 Hz), high beta (18–30 Hz), and beta (12–30 Hz) of each brain region were derived by Fast Fourier Transformation (6). Two normalization strategies were applied to power analysis, subject- and frequency-wise, with the former normalized to the whole brain power and the latter to the summed power of each studied frequency band. The interaction between brain regions for each frequency band was calculated by lagged coherence and lagged phase synchronization (7). The equations for phase synchronization are the same as those for coherence except for a pre-normalization step to discount the influence of power, hence non-linear. In other words, the relationship between amplitudes did not affect phase synchronization. With the metrics of regional power and inter-regional connectivity established, we performed similar computations like the fMRI counterpart—correlation between regional power and the mean connectivity strength around that region. In this exploratory analysis, we correlated the mean power and mean connectivity across all the subjects. The results showed an interesting pattern that at lower frequencies (delta and theta), there remained a robust negative relationship. At a higher frequency range (alpha to beta), the negative relationship disappeared after the correction of multiple comparisons. Together, the findings indicate that the information carried in a lower frequency range (fMRI and EEG) is more relevant to inhibitory interaction. The results are summarized in **Supp Table 3**.

**Supp Table 3**

Statistics of the correlation between power and connectivity strength across different spectra.

Spectra	delta	theta	alpha1	alpha2	alpha	low beta	high beta	beta

Subj-Coh	-0.44	-0.74	0.01	0.32	0.22	-0.15	-0.30	-0.21

*P* value	1.68E-4	5.96E-13	0.91	0.01	0.08	0.21	0.01	0.09

Freq-Phase	-0.55	-0.69	-0.01	0.33	0.22	-0.16	-0.31	-0.24

*P* value	1.04E-6	1.17E-10	0.91	0.01	0.07	0.18	0.01	0.05

Subj-Coh	-0.44	-0.74	0.00	0.31	0.20	-0.15	-0.30	-0.21

*P* value	1.83E-4	6.49E-13	0.98	0.01	0.09	0.21	0.01	0.08

Freq-Phase	-0.55	-0.68	-0.02	0.33	0.21	-0.16	-0.31	-0.24

*P* value	1.08E-6	1.36E-10	0.86	0.01	0.08	0.19	0.01	0.05

Note: Subj and Freq represent two normalization strategies of power, i.e., subject- and frequency-wise. Coh and Phase represent two functional connectivity measures, i.e., lagged coherence and phase synchronization. The data are organized by paired rows, upper the correlation coefficients and lower the P values (N=47).
